# Integrated PET and confocal imaging informs a functional timeline for the dynamic process of vascular reconnection during grafting

**DOI:** 10.1101/2022.10.27.513862

**Authors:** Margaret H. Frank, Sergey Komarov, Qiang Wang, Ke Li, Matthew Hecking, Halley Fowler, Claire Ravenburg, Audrey Widmier, Arielle Johnson, Hannah Thomas, Viktoriya Coneva, Daniel H. Chitwood, Yuan-Chuan Tai

## Abstract

Grafting is a widely used agricultural technique that involves the physical joining of separate plant parts so they form a unified vascular system, enabling beneficial traits from independent genotypes to be captured in a single plant. This simple, yet powerful tool has been used for thousands of years to improve abiotic and biotic stress tolerance, enhance yield, and alter plant architecture in diverse crop systems. Despite the global importance and ancient history of grafting, our understanding of the fundamental biological processes that make this technique successful remains limited, making it difficult to efficiently expand on new genotypic graft combinations. One of the key determinants of successful grafting is the formation of the graft junction, an anatomically unique region where xylem and phloem strands connect between newly joined plant parts to form a unified vascular system. Here, we use an integrated imaging approach to establish a spatiotemporal framework for graft junction formation in the model crop *Solanum lycopersicum* (tomato), a plant that is commonly grafted worldwide to boost yield and improve abiotic and biotic stress resistance. By combining Positron Emission Tomography (PET), a technique that enables the spatio-temporal tracking of radiolabeled molecules, with high-resolution laser scanning confocal microscopy (LSCM), we are able to merge detailed, anatomical differentiation of the graft junction with a quantitative timeline for when xylem and phloem connections are functionally re-established. In this timeline, we identify a 72-hour window when anatomically connected xylem and phloem strands regain functional capacity, with phloem restoration typically preceding xylem restoration by about 24-hours. Furthermore, we identify heterogeneity in this developmental and physiological timeline that corresponds with microvariability in the physical contact between newly joined rootstock-scion tissues. Our integration of PET and confocal imaging technologies provides a spatio-temporal timeline that will enable future investigations into cellular and tissue patterning events that underlie successful versus failed vascular restoration across the graft junction.

## Introduction

Vascular plants have an innate capacity to regenerate in response to wounding, enabling them to be cut and grafted onto other individuals (1). This property was first recognized by ancient farmers thousands of years ago (2, 3), and has since become an essential technology that growers use to combine desirable traits between genotypically distinct root and shoot systems. This simple yet effective strategy is widely applied to vegetable and perennial crops to combat biotic and abiotic threats that would otherwise have devastating effects on crop production (4–6). Despite the tremendous advantages that grafting provides for crop protection and enhanced crop yield, successful graft combinations are often identified on an inefficient, case-by-case basis (7, 8). This is largely due to a present lack of understanding regarding the fundamental biological mechanisms that underlie successful graft pairings. In this study, we focus on a central determinant of a successful graft: the re-establishment of vascular function across the graft junction.

The graft junction is a unique anatomical region that unites newly joined root and shoot systems into a single vascular conduit. The temporal dynamics of graft junction formation varies widely among species. Studies from Arabidopsis indicate that phloem and xylem strands restore functional transport within 3-4 and 6-7 days post-grafting, respectively, under room temperature conditions (9), and that elevated temperatures accelerate this process by approximately 24 hours (10). Woody perennials, on the other hand, require anywhere from weeks to months to form stable junctions (11, 12). Regardless of whether the graft is formed between relatively fast-grafting, herbaceous species or slower woody perennials, there is a shared sequence of cellular events that ultimately leads to successful junction formation (13). Graft healing is initiated by cellular adhesion at the rootstock-scion interface, where a necrotic layer is formed from the wounded cells along the cut site. Following adhesion, necrotic clearing and proliferation of new callus parenchyma at the root-shoot interface fills in air spaces within the junction. Subsequent specification of callus into new vascular tissue leads to the formation of *de novo* xylem and phloem connections that bridge the rootstock and scion together (12). The graft junction is considered complete once these newly formed xylem and phloem strands mature into functional conducting tissue.

While substantial progress has been made towards identifying the molecular genetic processes that underlie the initial steps of graft adhesion and callus production (9, 15–17), relatively little is known about how vascular reconnection is coordinated within the developing junction. Basic questions such as how differentiation corresponds with functional re-establishment of physiological transport across the junction remain largely unexamined. These fundamental gaps in our understanding of how grafts are formed can in part be attributed to a lack of *in vivo* tools that allow for simultaneous quantitative measurements of xylem and phloem conductance. Currently, there are two methods that are used to investigate restored physiological transport during grafting. A common method for sensitive tracking of physiological transport involves the use of xylem- and phloem-mobile dyes and proteins (9, 18–20). Another, direct physiological measure looks at hydraulic conductance as a metric for restored xylem transport (21). Both of these methods are informative for tracking graft formation; however, they are also fundamentally limited. Dye transport assays typically require destructive harvesting, and thus are often not appropriate for tracking dynamic processes during graft formation. Moreover, it is challenging to simultaneously track xylem- and phloem-mobile dyes within the same individual, since loading the dye typically involves wounding, and there are a limited number of suitable dyes with non-overlapping emission spectra. Hydraulic conductance, on the other hand, can be performed on the same intact plant over multiple days, but it only provides a coarse measurement for xylem restoration, and cannot be used to track restored phloem function.

Positron emission tomography (PET) is a quantitative imaging technique that enables highly sensitive, non-invasive *in vivo* measurements of physiological processes (22–24). PET relies on the production of biomolecules that carry radioactive tags – typically radio-isotopes that decay through positron emission. These radioactive molecules, referred to as radiotracers, can be incorporated into living organisms to track metabolic and/or physiological processes in real time. As the radiotracers decay, they release positrons that react with nearby electrons, creating annihilation events that produce two gamma photons that are released near 180 degrees from one another. PET scanners contain multiple rings of detectors that can detect simultaneous photon release, allowing researchers to computationally reconstruct the position of radiotracers within living organisms in 3D over a course of minutes to days (depending on the type of radiotracers employed) (22, 23).

In this study, we develop a new method for tracking dynamic anatomical and physiological vascular restoration during graft formation. By integrating PET imaging with laser scanning confocal microscopy (LSCM), we are able to construct a high-resolution anatomical and physiological map for restored vascular function. The synthesis of these two datasets reveals a 72-hour window in which xylem and phloem strands differentiate and restore long-distance transport. Through this work, we also identify sources of micro-variability that alter the temporal progression of graft formation and should be considered as a potential factor impacting the molecular characterization of junction formation.

## Results and Discussion

### PET imaging enables dynamic tracking of xylem and phloem function

Restored physiological function following grafting has yet to be measured using non-destructive, *in vivo* approaches that allow xylem and phloem conduits to be tracked in the same individual. To address this technological gap, we tested radionuclides that could be delivered in a non-destructive manner, exhibit dynamic transport as xylem and phloem conduits, and have sufficiently short half-lives to allow for same-day dual labeling. To test whether we could directly track radiolabeled phloem conduits we delivered gaseous ^11^CO2 to ungrafted tomato seedlings using a chamber and an airtight seal around the base of the hypocotyl (Supplemental Figure S1). The labeled seedlings fixed the ^11^C radioisotope within 14 minutes post-labeling (Figure 1). Within 19 minutes post-labeling, the labeled photosynthates moved into the upper root system, and by 133-162 minutes post-labeling, ^11^C accumulated in the growing root tips of the seedling (Figure 1). To track dynamic xylem transport, we initially tested whether radiolabeled H_2_^15^O could be delivered to root systems and traced through the xylem; however, the short half-life for ^15^O (approximately 122 seconds) and high energy positrons emitted by this radioisotope produced a short-lived, signal with high levels of noisy background caused by positron escape (Supplemental Figure S2). We tested an alternative radiolabel, ^13^N, which has a half-life of approximately 10 minutes. By feeding plants water with labeled ammonium ([^13^NH4]^+^, referred to as ^13^N from here on out) we were able to directly deliver ^13^N to root systems; this produced an informative signal that moved from the labeled roots into the cotyledons within 17-minutes post-labeling (Figure 1B).

**Figure 1.**
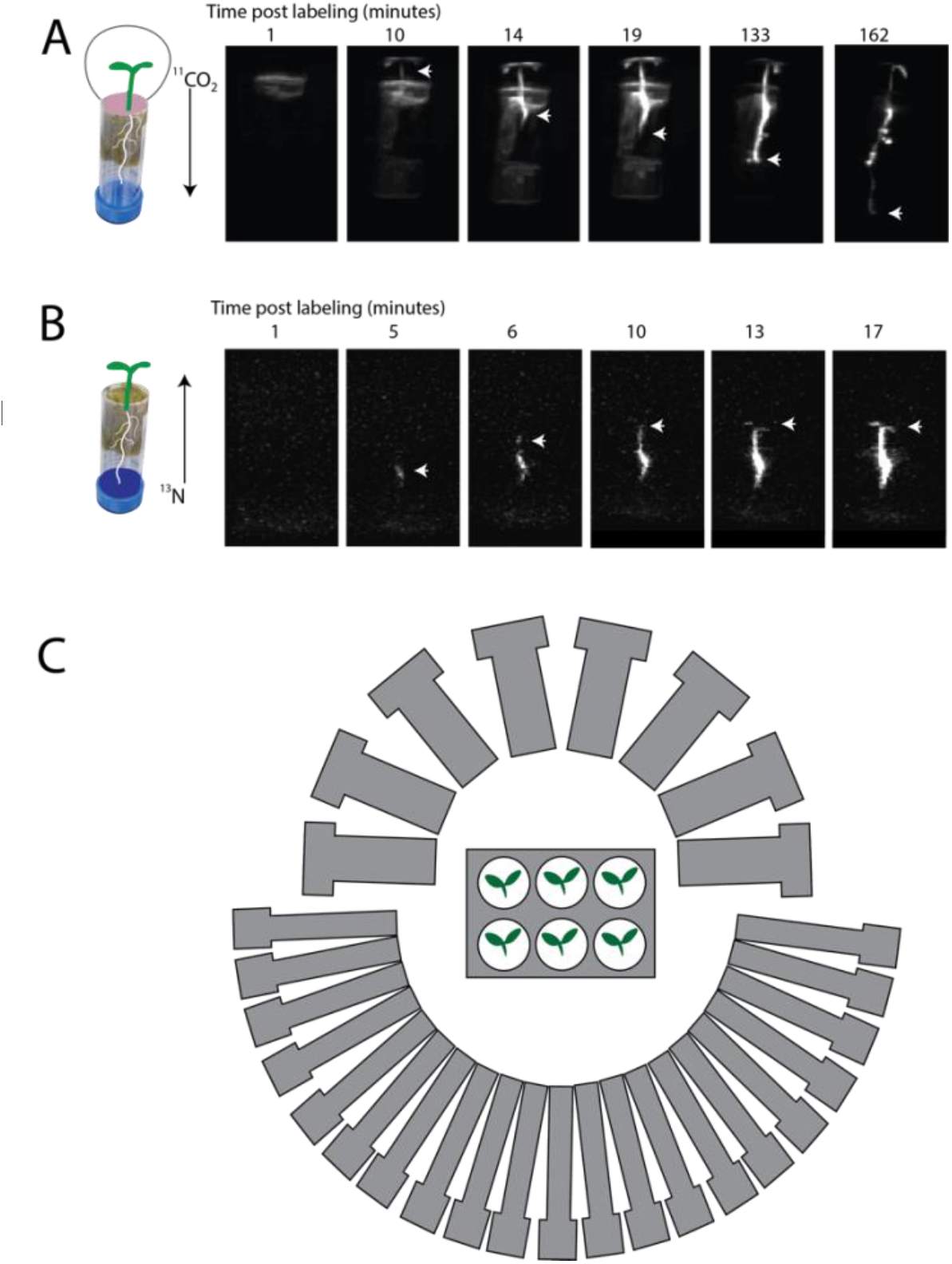
Dynamic PET imaging of phloem and xylem transport using ^11^CO_2_ (A) and ^13^N (B), respectively. Radiolabeled ^11^CO_2_ can be traced from young cotyledons where carbon is fixed into photosynthate, and transported through the phloem to the top of the root system within 14 minutes post-labeling (A). Distal root tips accumulate ^11^C-labeled photosynthate within less than 2.5 hours post-labeling. Xylem transport can be traced through the uptake and transport of ^13^N from seedling roots into cotyledons (B). Initial ^13^N uptake appears as a concentrated signal in the seedling roots within 5-minutes post-labeling, and moves acropetally into the cotyledons within 17 minutes post-labeling. The Plant PET imager is built inside of a Conviron growth chamber and consists of a large and a small half ring of 21 and 8 scintillation crystal detectors, respectively (23) (C). White arrows in (A & B) mark the moving front of radiolabeled ^11^C and ^13^C into the distal root and shoot systems, respectively. Dental impression media (yellow putty in A) was used to create an airtight seal above the graft junction region during ^11^CO_2_ labeling. Real pictures of the plants shown in Supplemental Figure S1.

Next, we used our PET labeling protocol to capture a quantitative timeline for the physiological restoration of xylem and phloem transport during graft healing. We performed several staggered grafting experiments that enabled us to simultaneously measure vascular transport across a range of developmental time points from 1-7 days post-grafting (Figure 2; Supplemental Figure S3). By including an ungrafted control in every imaging round, we were able to differentiate between technical failures while delivering the radionuclides and biologically informative failure to transport due to severed vascular connectivity across the junction (Figure 2). During the xylem transport experiments we discovered that ^13^N uptake is highly responsive to the water source that was used for our hydroponic growth conditions. We were able to remedy this issue by using distilled water with a neutral pH. This technical hurdle limited the replication of our xylem conductance measurements; nonetheless, we were still able to identify a time point for xylem restoration at 6 days post-grafting (Figure 2A).

**Figure 2.**
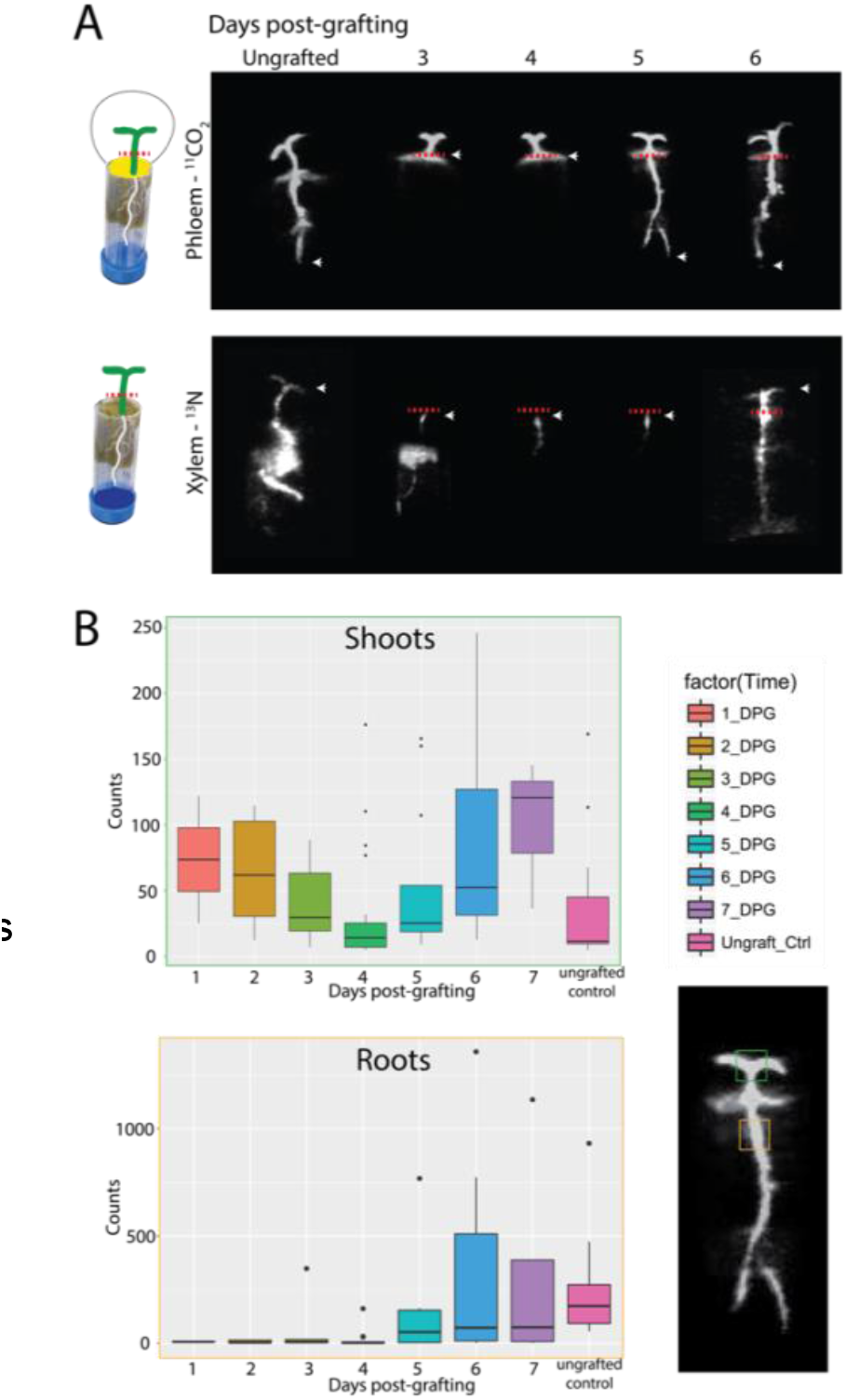
Physiological timeline for xylem and phloem restoration during graft junction formation delineates a window for vascular maturation. Radiolabeled ^11^C fed to grafted scions in the form of ^11^CO_2_ is transported to rootstocks through the phloem by 5 days post-grafting (A). Radiolabeled ^13^N fed to grafted rootstocks accumulates at the graft site in healing junctions 3-5 days post-grafting, and initiates root-to-shoot xylem transport into the scion by 6 days post-grafting (B). Region of interest quantification of radioactive accumulation in grafted scions and rootstocks demonstrates that phloem transport is restored by 5 days post-grafting (C). ^11^CO_2_ data was collected 7200-14400 seconds after labeling and ^13^N data was collected 3600 -7200 seconds after labeling. Red dotted lines in (A) indicate the position of the graft junction, and the green and orange boxes in (C) indicate scion and rootstock ROIs, respectively.

To supplement the reduced sampling for ^13^N xylem transport, we used a standard 5(-and-6)-Carboxyfluorescein Diacetate (CFDA) dye transport assay (9, 19) to track the flow of xylem-mobile fluorescent dye from severed roots into cotyledons. The CFDA transport assay aligned closely, but not perfectly, with our PET imaging results, showing stabilized xylem transport at 5 days post-grafting (DPG) (Supplemental Figure S4). Thirty percent of the plants showed xylem transport of CFDA as early as 4 DPG. To gain further insight into xylem formation at this early time point, we examined the anatomical connectivity of the graft junction using confocal microscopy and found that these grafts lacked mature vessel connections (Supplemental Figure S4). Prior to 6 DPG, our PET data showed ^13^N radioactivity pooling within the junction region (Figure 2). This can be attributed to positive xylem pressure that causes ^13^N containing xylem sap to exude out of unhealed vessel elements and into the apoplast of the junction. Nitrogen requires active transporters to re-enter the symplast, and thus becomes stuck at the graft site once exuded. Unlike ^13^N, CFDA can diffuse back across the junction and re-enter the vascular transport stream on the scion half of the graft. The small differences in temporal transport that we observed between our PET and CFDA assays is likely due to physiological differences between ^13^N and CFDA transport dynamics.

To determine a temporal timeline for restored phloem function, we waited for the ^13^N radionuclide to degrade over a period of more than 10 half-lives (> 100 minutes), and then labeled the same plants with gaseous ^11^CO_2_. We used airtight labeling chambers and dental impression putty that blocked ^11^CO_2_ from being delivered below the graft junction (Figure 2; Supplemental Figure S1). To measure restored phloem transport, we quantified the intensity of radioactivity within a region of interest (ROI) above and below the graft junction (Figure 2B). In unhealed junctions from 1-4 DPG, we noticed that shoot-to-root phloem transport was abruptly halted at the site of the graft junction, and remained within the scion for the duration of our data collection (≥ 180 minutes post-labeling) (Figure 2A). By 5 DPG, we observed restored phloem transport in 50% of the grafted seedlings (Figure 2B; Supplemental Figure S3). Subsequent healing from 6-7 DPG led to plants with increased radioactive accumulation within the rootstock, indicating increased transport capacity within the developing phloem (Figure 2B).

### Vascular maturation follows a 72-hour timeline from anatomical initiation to physiological restoration

To examine anatomical restoration during graft formation, we stained and optically sectioned junctions from 3-9 DPG using laser scanning confocal microscopy (LSCM). As early as 3 DPG we observed initial signs of vascular differentiation in both well-formed (Figure 3A) and slightly off-center grafts (Figure 3B). At this stage of regeneration, we observed cells along the graft site with isodiametric callus-like morphology that also formed annular secondary cell wall thickenings that are indicative of protoxylem identity, thus forming an intermediate callus-protoxylem cell type (Figure 3A-C). In contrast, junctions with incomplete cellular contact across the graft site formed protruding callus cells with protoxylem-like secondary cell wall thickenings and air pockets within the internal cellular structure of the junction (Figure 3B-C). At the 4 DPG timepoint, we could distinguish immature protoxylem strands that connected across the junction (Figure 3E-G). These vascular strands often contained bulbous, callus-shaped vessel elements (prominently shown in Figure 3), bordered by tapered phloem tissue along the periphery of the developing junction (Figure 3H).

**Figure 3.**
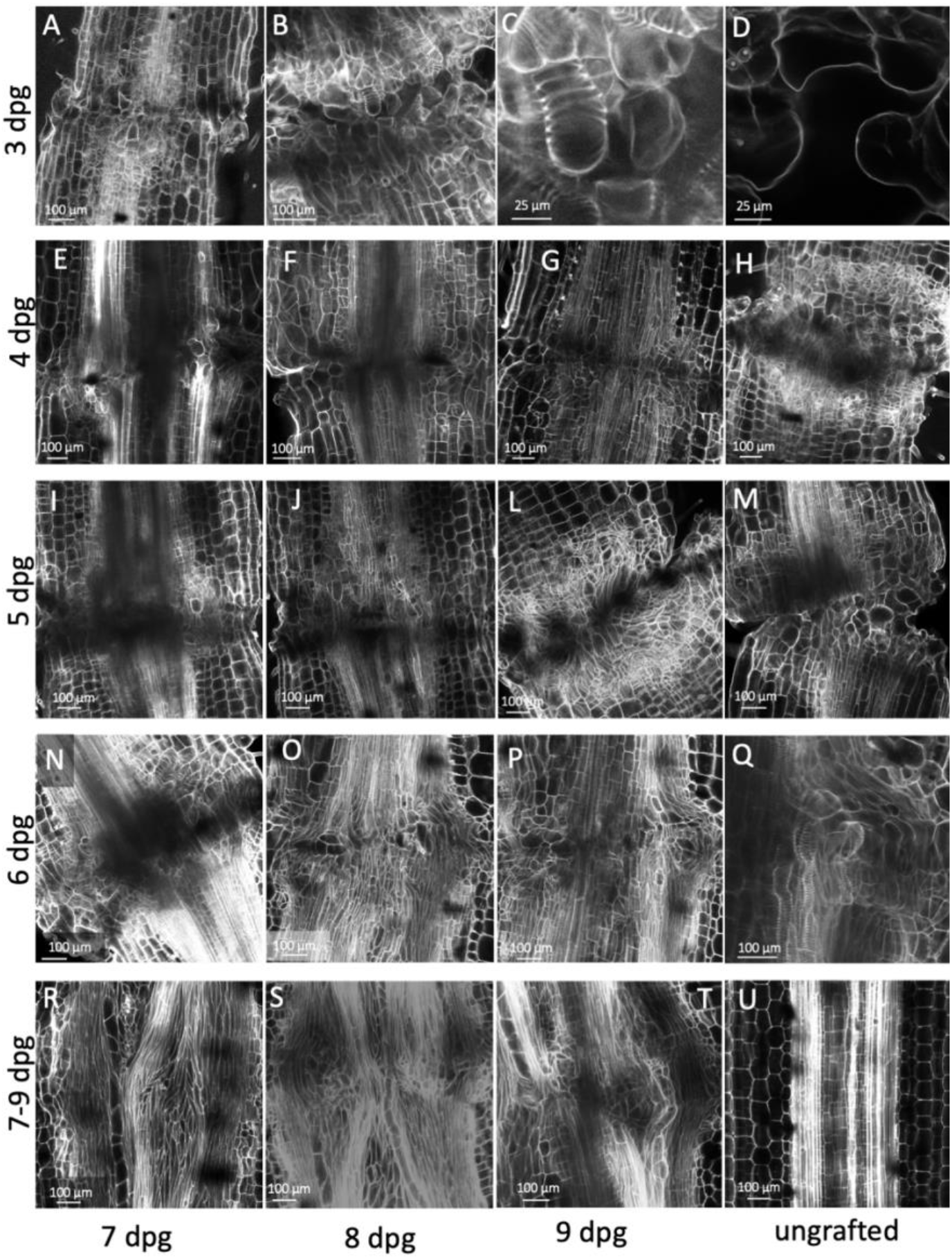
Anatomical differentiation of the graft junction region from 3-9 days post-grafting defines a window for vascular differentiation and sources of developmental variability. Optical sectioning using confocal microscopy of propidium iodide stained junctions was used to construct an anatomical timeline for junction formation from 3-9 days post-grafting. A well connected graft junction imaged at 3 days post-grafting (DPG) shows stabilized callus connections between the rootstock and scion with intermediate callus-protoxylem cells that have isodiametric morphology characteristic of calli and annular secondary cell wall thickenings that mark protoxylem identity (A). Grafts with poor rootstock scion contact at 3 DPG (B) form callus and hybrid callus-protoxylem cellular projections from both sides of the junction that fail to make contact across the graft (close up of B shown in C). Air spaces are visible within grafts that form poor cellular contact across the junction (D). Immature xylem strands can be traced across graft junctions by 4 DPG (E-H); however, most of these strands include bulbous protoxylem cells that have not matured into conducting tissue (indicated with arrowheads in E & G). Phloem strands can also be visualized connecting across the graft junction along the periphery of newly formed xylem strands (Labeled in a median section through the junction in E, and a tangential section through differentiating phloem in H). By 5 DPG well-aligned grafts have formed stabilized xylem and phloem connections across the junction (I-L) while grafts with poor rootstock-scion contact are anatomically delayed and developmentally resemble junctions at the 3 DPG stage (M). By 6 DPG, mature vessel elements and sieve tubes are connected across the junction, which coincides with restored xylem and phloem transport (N-P, and close up of connected xylem and phloem strands from P shown in Q). Extensive vascular differentiation from 7-9 DPG produces junctions that are packed with mature metaxylem, protoxylem, and phloem connections, and lack pith cells (R-T). In contrast, ungrafted hypocotyls have parallel vascular strands that surround a central pith (U).

We could clearly identify stabilized vascular bridges curving through the junction by 5 DPG (Figure 3I-L). While the protoxylem bridges could easily be traced across the junction at this timepoint, we noticed that the vessel elements comprising these bridges presented a spectrum of cellular morphologies from bulbous to elongated shapes, corresponding with differing degrees of vascular maturation. This morphological spectrum aligned closely with the functional window that we observed for physiological xylem restoration in our PET and CFDA xylem transport experiments (Figure 2A; Supplemental Figure S4). We also observed off-centered grafts at 5 DPG that showed substantial delays in vascular differentiation, anatomically resembling well-connected grafts at 3 DPG (Figure 3M). We suspect that this delayed vascular differentiation in off-centered grafts is caused by a prolonged callus proliferation phase, and that these micro-fluctuations in rootstock-scion alignment likely explain the temporal variation in physiological restoration that we observed with our PET imaging (Figure 2).

By 6 DPG, our PET data demonstrates that xylem and phloem transport is fully restored (Figure 2). In association with this physiological milestone, we could distinguish mature protoxylem, metaxylem, and phloem strands connecting rootstocks and scions (Figure 3N-Q). From 7-9 DPG, we observed continued maturation of surrounding xylem tissues that weaved through the junction, disrupting the central pith (Figure 3R-T). Another pronounced anatomical feature that we saw at 1 week post-grafting was the presence of wide metaxylem strands that curved through the regenerated graft region marking the re-establishment of efficient root-to-shoot transport (Figure 3N-Q). Overall, our anatomical imaging aligns closely with a model in which physiological transport does not resume until reprogrammed vascular tissue differentiates into mature, elongated cells that span the junction at 5-6 DPG (Figure 4).

**Figure 4.**
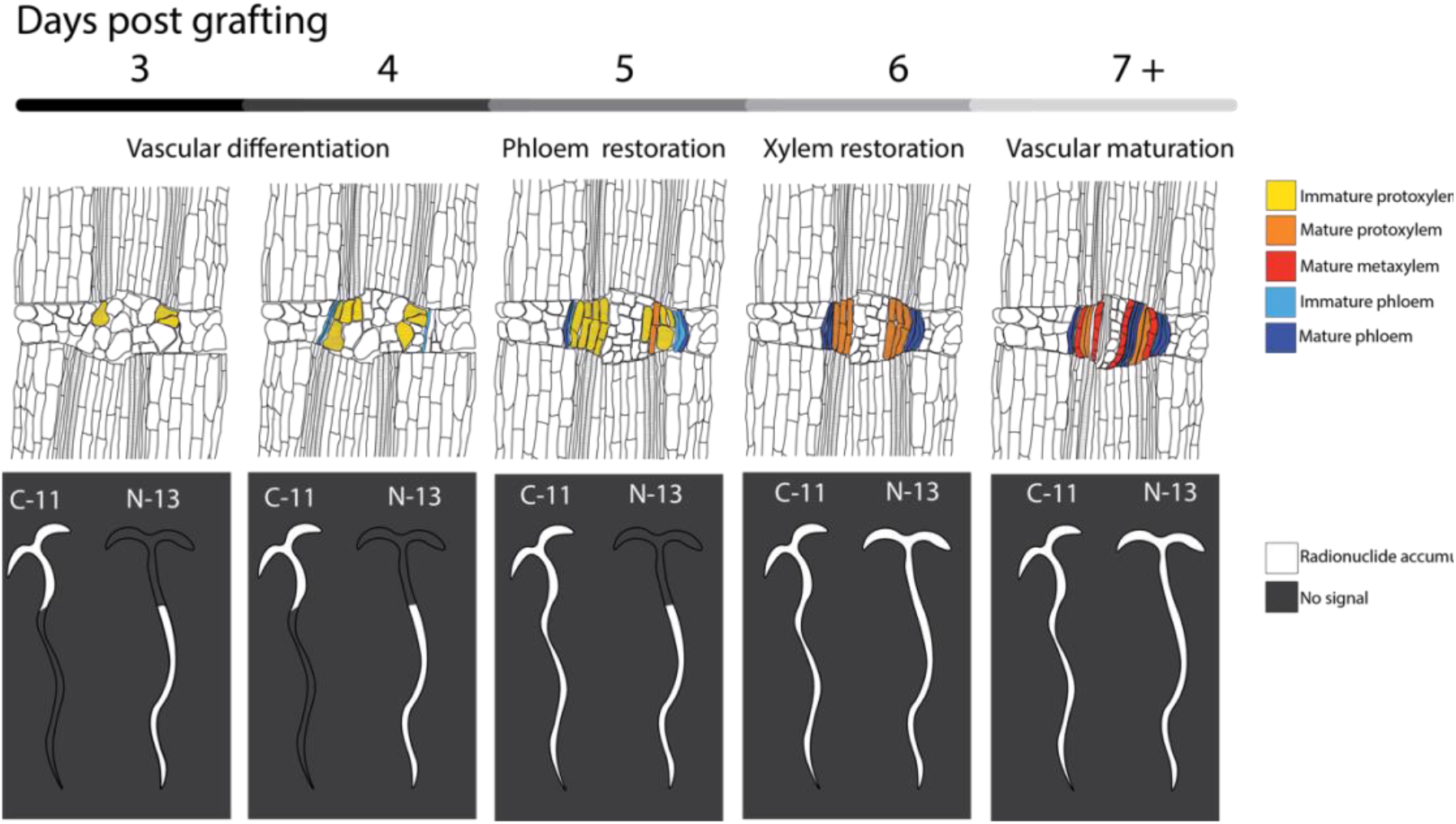
A model for the anatomical and functional dynamics of graft junction formation informed by PET and confocal imaging. Within 3 days post-grafting (DPG), early signs of *de novo* protoxylem specification are present. These premature protoxylem cells initiate with a callus-like morphology and subsequently differentiate into elongated vessel elements between 4-6 days post grafting, corresponding with restored root-to-shoot xylem transport. Phloem strands are visible by 4 DPG and subsequently show extensive connectivity between 5-6 DPG, corresponding with restored shoot-to-root phloem transport at 5 DPG.

## Conclusion

We show that PET imaging can be used to sensitively track *in vivo* physiological function during graft regeneration. By pairing PET and LSCM technologies together, we are able to access fundamental questions regarding the relationship between anatomical maturation and restored transport. Our work identifies a 3 day developmental window between the first anatomical signs of vascular differentiation to restored physiological transport (Figure 4). Furthermore, we show that vascular transport does not resume until differentiating vasculature reaches maturity. Finally, we identify sources of heterogeneity during junction formation that not only impact developmental staging during molecular investigations into graft formation, but may also influence anatomical connectivity and ultimately the success of agricultural grafts.

## >Materials and Methods

### Growth conditions and plant grafting

*Solanum lycopersicum* Cv. M82 seeds were treated with 50% Bleach for 30 seconds, rinsed thoroughly with deionized water (diH_2_O), and placed on damp paper towels to germinate in Phytatrays (Sigma-Aldrich, Saint Louis, MO, USA). Germination was synchronized by giving the seeds 3 days of dark treatment at room temperature, followed by three days in the growth chamber. The seedlings were then transferred into bottomless 50 mL Falcon tubes lined with rockwool (Grodan, Roermond, Netherlands), and grown at 23 ºC, with a repeating 16:8 light:dark cycle in a hydroponic tub supplied with aeration rocks. Custom grafting clips that securely hold the rootstock and scion together at this early seedling stage were made by cutting a slit down the side of Creatology Round Plastic Lacing (Michael’s Craft Store, Irving, TX, USA). Scions between neighboring seedlings were reciprocally grafted onto one another using the tube grafting method on a staggered timescale starting 24 hours after plants were transferred into the hydroponic tub. Unsuccessful grafts were identified based on the appearance of drooping cotyledons within 48 hours post-grafting and were removed from the study. Notably, some grafts died mid-study and were pulled from the dataset.

### Delivery and PET imaging of ^13^N and ^11^C radionuclides

*In vivo* functional xylem and phloem reconnections were investigated using PlantPET, a PET scanner with a vertical bore inside a plant growth chamber dedicated to functional plant imaging research (23) (Figure 1; Supplemental Figure S1). Functional xylem transport was assessed by feeding the rootstocks trace concentrations of aqueous radiolabeled ^13^N in the form of ammonium (Fig 2). The choice of aqueous ^13^N-labeled tracer over ^15^O-labeled water (H_2_^15^O) to measure flow in the xylem was based on: (1) the relatively short half-life of ^15^O (2 minutes), which limits the amount of time available for imaging, resulting in noisy images; and (2) the fact that ^15^O emits high energy positrons that have a higher probability to escape the small tomato plants before positron annihilation, which increases the noise outside of the subject and reduces the usable events within the stem and leaf (Supplemental Figure S2). 1 mCi of [^13^NH4]^+^ dissolved in water was delivered to the bottom of the 50 mL Falcon tube using a syringe (illustrated in Fig. 1B). The tubes were placed in a custom-built lead rack that holds six plants (Figure 1; Supplemental Figure S1) and cuts down background noise by shielding the scanner from radioactivity in the Falcon tubes. These plants were placed into the PlantPET imager where they were imaged for 10 minutes to gauge radioactivity uptake. The radioactivity was flushed from the root system 30 minutes after labeling and replaced with diH_2_O, and then placed back in the PET imager for a continuous hour of imaging. Following >10 half-lives of ^13^N decay (>100 minutes), the plants were removed from the PET imager and prepped for ^11^CO_2_ delivery.

Functional phloem transport was assessed by feeding the leaves and shoots tracer concentrations of gaseous ^11^CO_2_ for photo-assimilation. In order to ensure that ^11^C was only fixed by portions of the plant that are above the graft junction, an airtight seal starting above the graft junction and extending to the edges of the Falcon tube was made out of Cinch dental impression media (Parkell, Brentwood, NY, USA) (Fig 2A). The plants were then encased in custom-made airtight labeling chambers, and fed 1 mCi of ^11^CO_2_. Following labeling, the plants were placed in the PlantPET imager where they were imaged continuously for 2 hours starting at approximately 20 minutes post radiolabeling.

### PET image reconstruction, signal quantification, and decay time corrections

Radioactive decays registered by the PlantPET scanner are stored as a continuous stream of list-mode events that can be divided into an arbitrary number of frames chosen by the user. We divided both sets of list-mode data (1-hour from ^13^N experiment and 2-hours from ^11^C experiment) into multiple subsets that are equivalent to a 60-second frame at the beginning of the imaging session. The actual frame duration was increased at later time points to account for radioactive decay (T_1/2_ = 9.97 min for ^13^N and 20.38 min for ^11^C). The same imaging protocol and frame definition was used for all xylem and phloem measurements (i.e., one for ^11^C and one for ^13^N experiments).

List-mode events in each of the time frames were reconstructed using a graphic processing unit (GPU) based fast list-mode image reconstruction algorithm (see (23)). The result is a time series of 3D image volumes (256×256×160 voxels of 0.8×0.8×0.8 mm each) that represent the distribution of radioactivity concentration in the plants over time.

### Rendering visual PET data and quantifying regions of interest (ROI) in FIJI

Reconstructed files were rendered as 3-D movies using the import raw stack and 3-D reconstruct functions in FIJI (25). ^11^C uptake and transport was quantified in FIJI using the ROI calculator function. A 7×7 pixel region of interest (ROI) was selected above and below the graft junction and average signal intensity across the 3-dimensional stack was quantified with the ROI calculator in FIJI ROI. Slices that dropped below an intensity value of two were considered to be products of “positron escape” and were removed from averaging. Transport was plotted as relative signal intensity above and below the graft junction. A similar calculation for ^13^N was not performed due to the non-biological background signal produced by the escaped positrons that annihilated in the Rockwool.

### Optical sectioning of the graft junction using laser scanning confocal microscopy

Graft junctions were harvested from 3-8 days post-grafting and vacuum infiltrated with ice-cold FAA (4% formaldehyde, 5% glacial acetic acid, 50% ethanol, 35% milliQ H_2_O). Samples were fixed overnight at 4 ºC, dehydrated through an ethanol series, rehydrated, stained for 1 hour in 0.002% propidium iodide (Thermo Fisher, Waltham, MA, USA), dehydrated again, and cleared for at least 10 days in 100% methyl salicylate (Sigma-Aldrich, Saint Louis, MO, USA)(26). The stained and cleared junctions were imaged on a Leica SP8 laser scanning confocal microscope with a white light laser set to 514-nm wavelength, with the laser intensity ranging from 10%-80% depending on sample depth. Junctions were optically sectioned using the Z-stack function on the LasX Leica Software, with a step size ranging from 1-1.4 µm per slice. Raw confocal images were imported into FIJI (25) and exported as single-frame images and multi-frame videos. Anatomical progression of graft junction vascular differentiation was analyzed by tracking changes in cellular morphology, secondary cell wall features that are characteristic of protoxylem and metaxylem cell identity, and the spatial arrangement of newly differentiated vascular strands within the junction.

### Xylem transport assays using CFDA dye

Replicate trials of soil grown grafted plants from 3-7 days post-grafting were cut at their root tips and dipped into (5 mg/mL) of 5(-and-6)-Carboxyfluorescein Diacetate (CFDA) (Invitrogen®, Waltham, MA, USA). Plants were incubated under full-spectrum LED lights (Cirrus LED Grow Lights, Saco, ME, USA) for 60-90 minutes, and then scored for the presence or absence of CFDA in cotyledons using a Leica M205 dissection microscope (Leica Microsystems Inc., Buffalo Grove, IL, USA). Ungrafted seedlings were included as positive controls for each round of imaging.

## Acknowledgments

This work was supported by a grant to MHF from the MO/AR EPSCoR Plant Imaging Consortium (IIA-1430427/IIA-1430428), NSF REPS funding for AW (IOS-1942437). NYSTEM C029155 and NIH S10OD018516 awards to Cornell University’s Biological Resource Center to fund the u880 laser scanning confocal microscope. CR was funded by a Cornell Presidential Life Sciences Fellowship, and NSF DBI-1040498 a Major Research Instrumentation grant for the development of the PlantPET scanner.

## Supplemental Figures

**Supplemental Figure S1.**
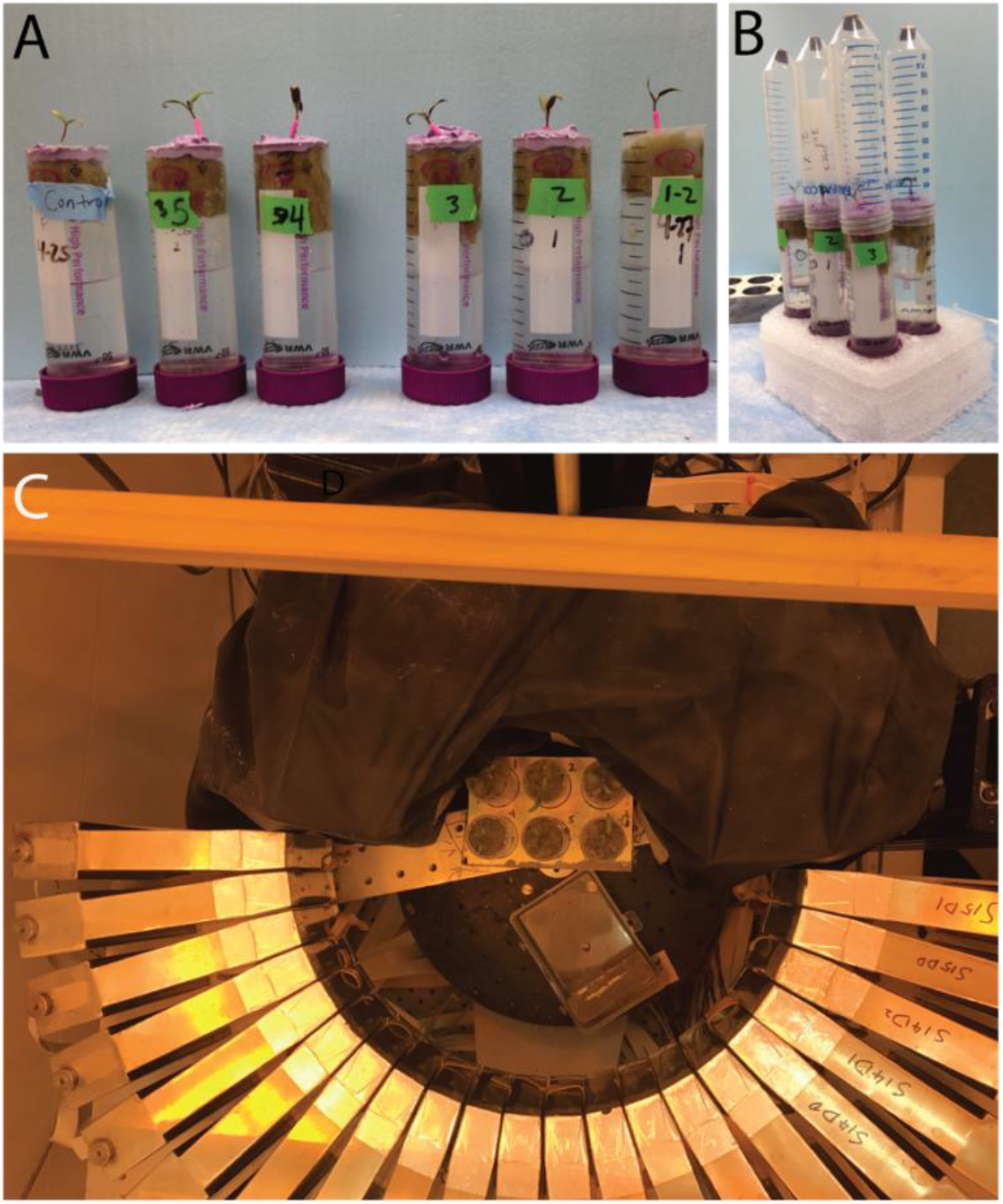
Plant labeling and imaging design. Grafted and ungrafted control plants were grown in rockwool plugs within 50 mL tubes. ^13^N was delivered to root systems (A) and gaseous ^11^CO_2_ was delivered to an airtight chamber encasing seedling shoots (B). Prior to labeling, plant tissue below the graft junction was sealed off from the air using dental impression media (shown in A and B). Plants were placed in a lead rack and positioned in the middle of the PlantPET imager for data collection (C).

**Supplemental Figure 2.**
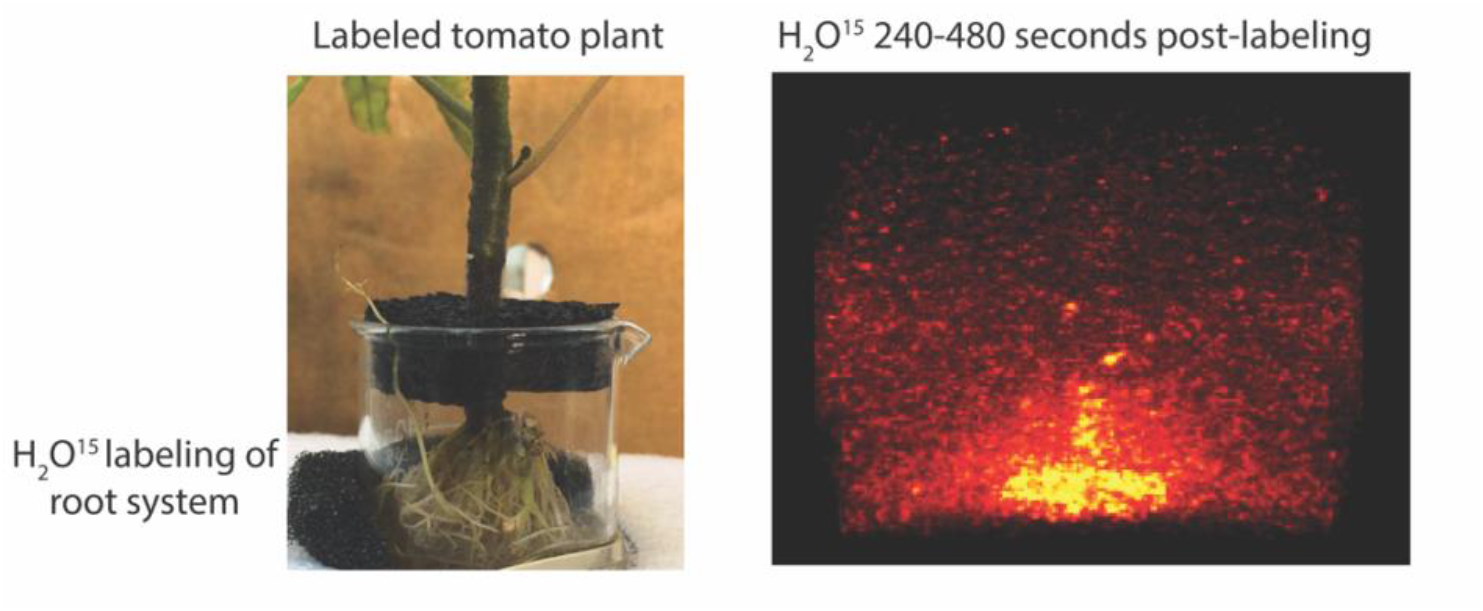
PET imaging of H_2_O^15^ to test whether O-15 could be used as a radiotracer. H_2_O^15^ has a half-life of 120 seconds, producing noisy PET imaging. Initial tests to track xylem transport using radiolabeled water failed due to the very short half-life of O^15^, and the high-energy positrons that this radioisotope emits, producing noisy data with a high level of background due to positron escape.

**Supplemental Figure 3.**
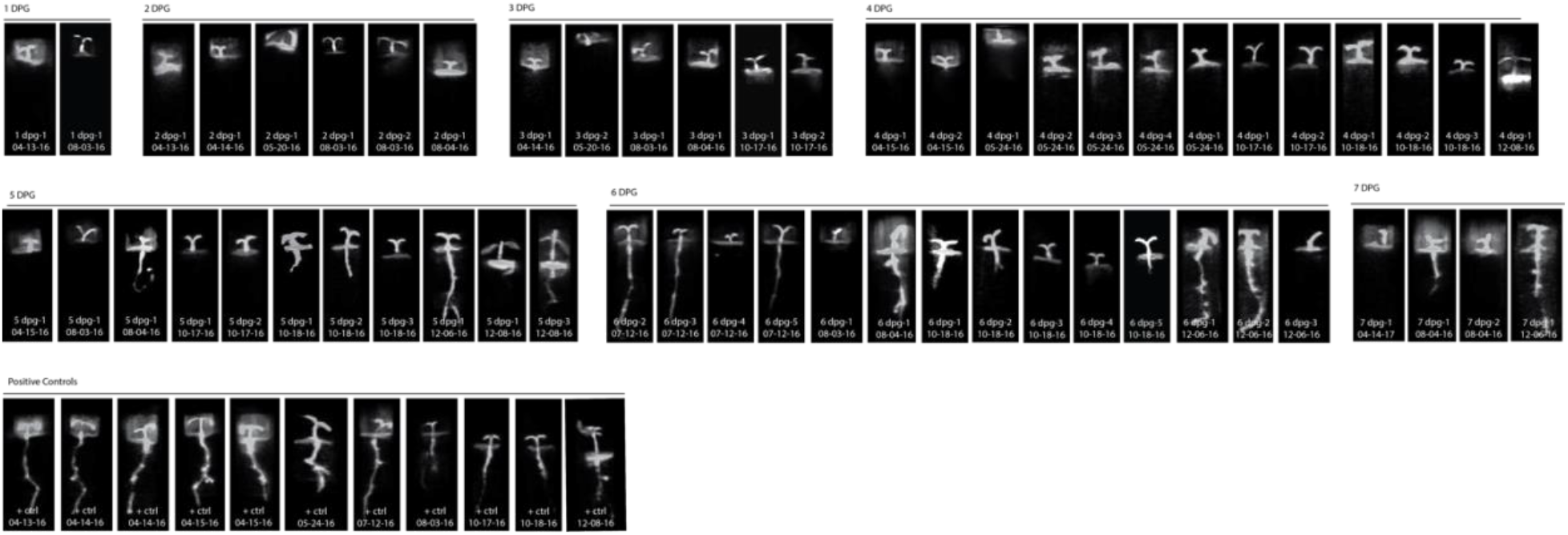
Complete images of ^11^C phloem transport subjects shows restored transport at 5 days post-grafting. Complete dataset of ^11^CO_2_ transport from 1-7 DPG, collected from 7200-14400 seconds post-labeling. Date and subject number are labeled at the bottom of each subject. Radiolabel accumulation corresponds with grayscale brightness in each image, with black indicating no radiolabel accumulation and white indicating the highest level of radioactive accumulation.

**Supplemental Figure 4.**
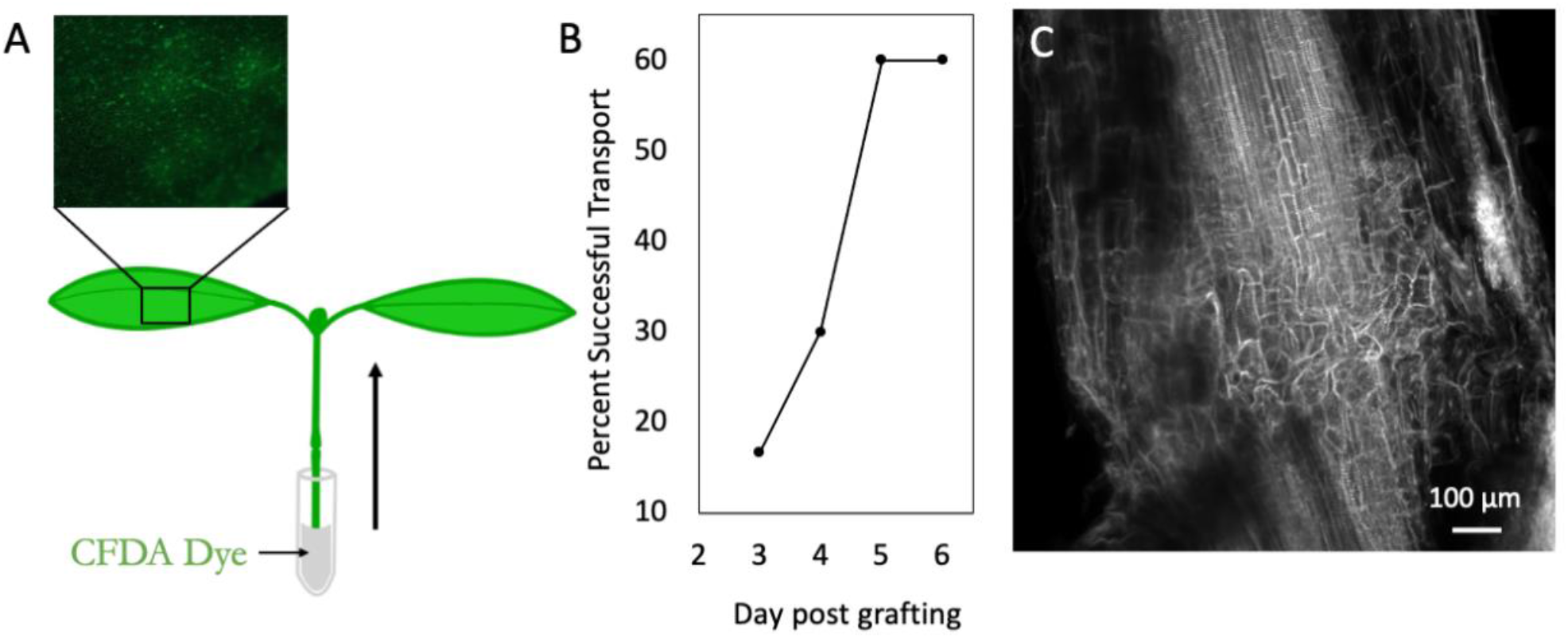
CFDA Dye transport assay for xylem restoration. Schematic for the 5(-and-6)-Carboxyfluorescein Diacetate (CFDA) transport assay to measure xylem transport in plants from 2-6 days post-grafting (A). 30% and 60% of grafts exhibited xylem transport by 4 and 5 days post-grafting, respectively (B). Max projection of confocal stack taken through graft junction at 4 days post-grafting shows that xylem files have not reconnected by this timepoint (C), indicating that low levels of dye transport can occur prior to xylem reconnection. Scale bar in C = 100 µm.

## Notes

### Competing Interest Statement

The authors have declared no competing interest.

